# Different patterns of eye-specific input are relayed to overlapping populations in the mouse superior colliculus

**DOI:** 10.64898/2026.09.22.753580

**Authors:** Miguel Ezeiza-Ortega, Jasmine Saluja, Jason W. Triplett

## Abstract

The mouse superior colliculus (SC) regulates critical behaviors such as prey capture and visual threat avoidance, each of which depend on binocular visual processing. In contrast to our previous understanding, recent work has demonstrated widespread and diverse binocular interactions in the mouse SC. The SC receives direct input from contralaterally and ipsilaterally projecting retinal ganglion cells (contra-and ipsi-RGCs), but the ways in which eye-specific information is filtered in the SC remain unclear. Here, we utilized a trans-synaptic, intersectional viral tracing strategy to label SC neurons receiving input monocularly from contra-RGCs (SC-C), monocularly from ipsi-RGCs (SC-I), or binocularly from both (SC-B). We show that a surprising number of SC neurons receive ipsi-RGC input and that the proportions of SC-C, SC-I, and SC-B neurons are consistent throughout the SC. Morphometric analysis revealed few differences between neurons receiving different eye-specific patterns of input. Consistent with this, we show that SC-C, SC-I, and SC-B neurons are comprised of similar proportions of five morphologically distinct subtypes. And, the expression of candidate molecular markers appears similar across innervation patterns. Together, these data suggest that eye-specific information is widespread throughout the SC targeting overlapping populations of neurons.

## INTRODUCTION

Binocular vision confers several advantages to species, including redundancy to unilateral damage, improved sensitivity, and depth perception^1^. In mice, binocular vision is critical for ethologically relevant behaviors, such as prey capture and visual threat avoidance^2,3^. The superior colliculus (SC) is a midbrain structure that processes multisensory information to regulate a wide range of behaviors^4^. Among these are prey capture and visual threat avoidance, which are mediated through distinct circuits downstream of the SC^5–7^. However, the ways in which binocular information is filtered through the SC to inform these processes remains poorly understood.

The SC receives direct binocular input from retinal ganglion cells (RGCs) that arise from both the contralateral and ipsilateral hemispheres (contra-and ipsi-RGCs). Across species, contra-and ipsi-RGC terminals are segregated into distinct patches or layers^8^. In rodents, the vast majority of retinal input to the SC is from contra-RGCs, which comprise ∼95% of the population and innervate the superficial-most layers of the SC. Ventral to this, ipsi-RGCs innervate the SC in a segregated layer where they terminate in patches localized primarily along the anterior and medial borders. The paucity of ipsi-RGC innervation and eye-specific segregation was thought to preclude binocular responses, and initial studies confirmed this^9–11^. In contrast to this, more recent work suggests that binocular interactions in the mouse SC may be more prevalent and diverse than previously thought^12–14^. Indeed, ∼60% of neurons in the mouse SC exhibit some form of binocular modulation. Beyond simple additive interactions in neurons that respond to stimulation through each eye individually, complex responses including facilitation, obligation, and suppression have been uncovered. Yet, how eye-specific information is distributed to give rise to such interactions remains unclear.

Furthermore, it remains unclear how different binocular modulation properties map onto neuronal subtype populations defined by visual tuning properties, gene expression, or morphology. Recent studies have begun to uncover the diversity of neuronal subtypes in the SC, though a unified framework has yet to be established. For instance, studies utilizing morphological criteria have identified 4-6 different subtypes^15–17^, while those leveraging gene expression criteria reveal 10-30 subtypes^18–20^. And, more discrete subtypes can be identified functionally, depending on the choice of visual stimulus^21–23^. Understanding how each of these subtype-defining criteria are inter-related will provide insight into how the SC processes and prioritizes visual cues to inform behavioral decisions.

We posit three models to explain how eye-specific information arriving in the SC could be relayed, each conferring distinct advantages. In a parallel model, eye-specific innervation patterns could be dedicated to distinct channels, each targeting specific subsets of SC neuron subtypes. An advantage of such organization could be efficiency of processing to mediate ethologically critical behaviors. In a distributed model, each subtype of SC neuron could receive all possible combinations of eye-specific relay. Such a strategy could have the advantage of increasing the feature space encoded across SC subtypes, allowing for more flexible visuomotor transformations. A third possibility is a mixed model in which some eye-specific innervation patterns are dedicated to distinct subtypes, while others are distributed across all subtypes.

To begin to test these models, here we leveraged a trans-synaptic and intersectional viral tracing method to label SC neurons receiving input monocularly from contra-RGCs (SC-C), monocularly from ipsi-RGCs (SC-I), or binocularly from both (SC-B). Surprisingly, while a majority of neurons are SC-C, a disproportionate number of neurons are SC-I, and a small population are SC-B. Interestingly, each type of innervation pattern was observed throughout the SC, in contrast to expectations based on the topographic localization of ipsi-RGC terminals. Morphological reconstructions of labeled neurons revealed that SC-I neurons have smaller somas than SC-C or SC-B neurons, but that other morphometric features were largely similar. Unsupervised clustering methods identified five morphologically distinct subtypes, which were also evenly distributed throughout the SC. Strikingly, similar proportions of morphologic subtypes were SC-C, SC-I, and SC-B and vice versa. Similarly, we observed no differences in the expression of candidate molecular markers between innervation patterns. Together, these data support a distributed model of eye-specific relay to the SC.

## MATERIALS & METHODS

### Mice

All experiments were performed in C57BL/6J mice obtained from Jackson Laboratories. Mice were housed in the Comparative Medicine Unit at Children’s National Hospital under standard conditions (12/12 light/dark cycle, *ad libitum* food and water). All experimental procedures were approved by the Children’s National Hospital’s Institute Animal Care and Use Committee.

### Intraocular injections

Mice aged postnatal day 30-31 (P30-31) were anesthetized with isoflurane and head-fixed in ear bars. A 30 gauge sharp tip needle was used to make a hole at the border of the cornea and sclera. A 30-gauge blunt needle attached to an Intraocular Injection syringe (Hamilton) was inserted into the posterior chamber and 1 mL of AAV1-hSyn-Cre-WPRE-hGH (AAV1-Cre, Addgene #105553) or AAV1-EF1a-FlpO (AAV1-FlpO, Addgene #55637) adeno-associated virus (AAV) was injected. Mice received subcutaneous injection of Buprenorphine (0.3 mg/kg) for post-operative analgesia.

### Stereotactic injections

Four days after intraocular injections, mice were anesthetized with isoflurane and placed in a stereotaxic apparatus. Focal craniotomies were made with a microdrill (Dremel) directly above both SC hemispheres (3.5 mm caudal and 0.4 mm lateral to bregma). A Hamilton syringe containing a 50/50 mixture of either AAV8-FLEX-tdTomato (Addgene #28306) and AAV9-Ef1a-fDIO-eYFP (Addgene #55641) or AAV9-pCAG-FLEX-eGFP (Addgene #51502) and AAV8-Ef1a-fDIO-mCherry (Addgene #114471) was lowered into the SC (1.5 mm ventral to bregma) and 1µL injected at a rate of 0.09 mL/min. After allowing 5 min following injection for viral diffusion, the needle was carefully withdrawn. Mice received subcutaneous injection of Buprenorphine for post-operative analgesia.

### Immunohistochemistry

Three weeks after stereotactic injections, mice were perfused intracardially with phosphate buffer solution (PBS), followed by 4% paraformaldehyde (PFA). Brains were extracted and post-fixed in 4% PFA overnight. Brains were then sectioned coronally at 150 mm intervals with a VS1000 Vibratome (Leica). Sections that contained the SC (∼3.08 – ∼4.72 mm caudal from bregma) were collected, maintaining their antero-posterior orientation. Tissue was cleared by CUBIC Clearing Protocol, as previously described^24^. Briefly, tissue was washed with CUBIC Reagent 1 overnight at room temperature, washed in PBS, and incubated in blocking solution (5% Donkey Serum, 0.5% Triton-X and 0.001% Na-azide in PBS) for 3 hours at room temperature. Sections were then incubated with primary antibodies (Table 1) diluted in blocking solution at 4 ℃ for 72 hours. Following this, sections were washed three times in PBS for one hour and incubated with secondary antibodies diluted in blocking solution at 4 ℃ for 48 hours. Sections were then washed three times in PBS for one hour and incubated with CUBIC Reagent 2 overnight at room temperature. Sections were mounted in CUBIC Reagent 2 using Frame Seal incubation chambers (Bio-Rad #SLF0601).

**Table 1.**
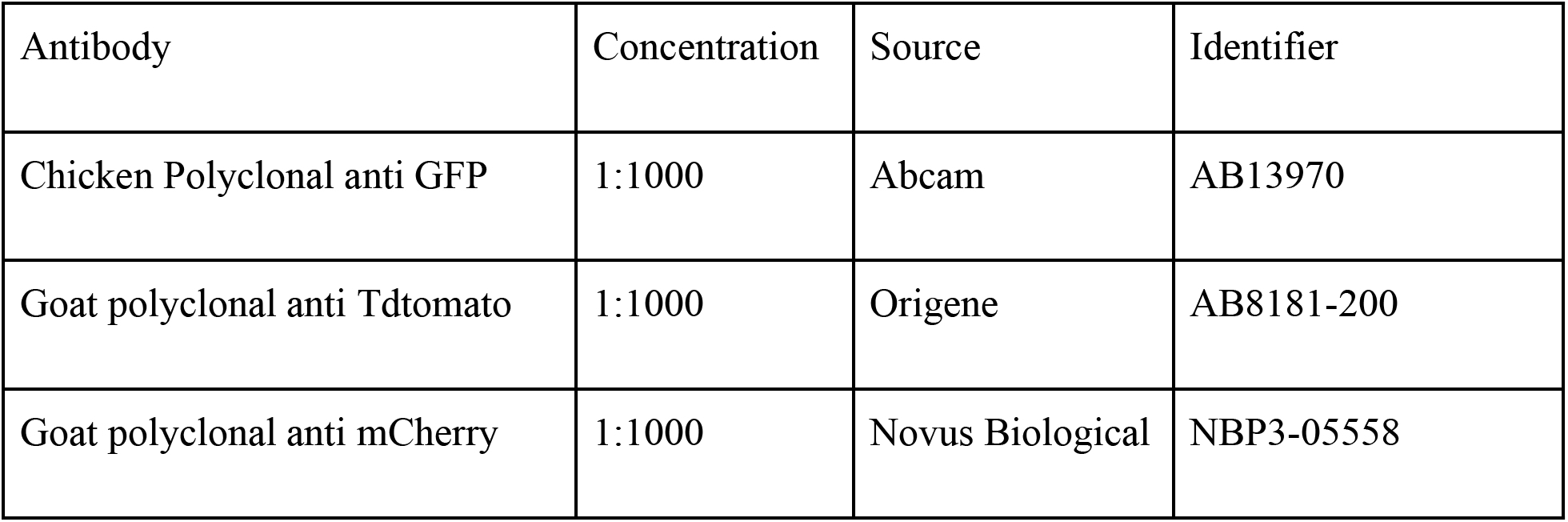
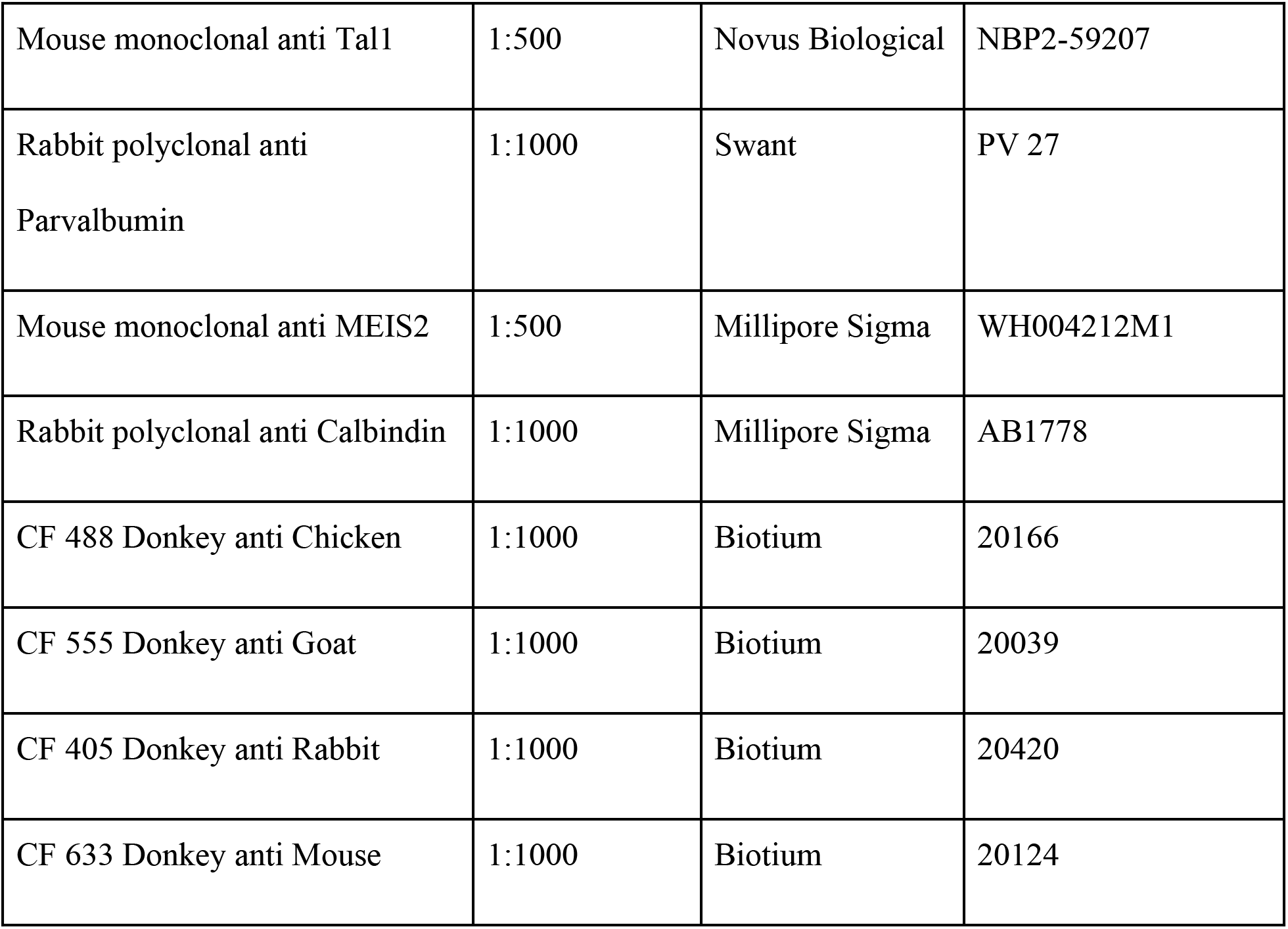
– Antibodies.

**Table 2.** – Viruses.

| Virus | Source | Identifier |
| --- | --- | --- |
| pENN-AAV1-hSyn-Cre-WPRE-hGH | Addgene | 105553 |
| pAAV1-EF1a-Flpo | Addgene | 55637 |
| AAV8-FLEX-tdtomato | Addgene | 28306 |
| AAV9-Ef1a-fDIO-eYFP | Addgene | 55641 |
| AAV9-pCAG-FLEX-eGFP | Addgene | 51502 |
| AAV8-Ef1a-fDIO-mCherry | Addgene | 114471 |

### Imaging and image analysis

Image stacks were obtained using a Nikon A1 confocal microscope at 40X magnification with a Z-resolution of 1 mm. Images were acquired from each SC hemisphere spanning the entire anterior-posterior axis, with roughly equal sampling along the medio-lateral axis. To prevent sampling bias, we attempted to count and reconstruct every neuron present in each Z-stack, though some were excluded due to their proximity to the edge of the section.

To identify somas labeled with each fluorescent protein and antibody, we utilized the “Surface” feature in Imaris (Oxford Instruments). To optimize thresholding, we delimited the foreground and background of 2-3 somas per fluorophore per image and automatically counted somas positive for each combination of fluorophore(s). Morphological cellular reconstructions were performed using the “Filament” feature in Imaris. Several morphometric features were extracted for each reconstructed neuron, including soma volume, total dendritic length, primary dendrite diameter, number and order of dendritic branches, and convex hull volume. In addition, Sholl intersections were counted at 5 mm radii to determine the total number of intersections, the maximum number of intersections, and the critical radius defined as the median radius at which the maximum occurred. In addition, we counted the number of Sholl intersections in four quadrants relative to the surface of the SC and used these data to determine the arbor height, arbor width, and arbor orientation index (AOI). As described previously^17^, AOI was calculated as *I_V_*⁄*I_H_*, where *I_V_* and *I_H_* are the number of intersections in vertical (dorsal and ventral) and horizontal (lateral and medial) quadrants, respectively. We also calculated the Sholl Ramification Index (SRI) as *I_max_*⁄*B*_1_, where *I_max_* is the maximum number of Sholl intersections and *B_1_* is the number of primary branches, and the Dendritic Branch Order Index (DBOI) as 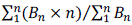, where *B* is the number of branches and *n* is the branch order.

### Clustering

We performed unsupervised *k*-means and hierarchical clustering based on morphological features to identify different cell classes of labeled neurons, as described previously^12,25^. The following 10 parameters were utilized for clustering, number of primary dendrites, average primary dendrite diameter, maximum Sholl intersections, total Sholl intersections, critical radius, arbor width, arbor height, AOI, DBOI, and SRI. We determined the principal component utilizing the *pca* function in MATLAB and clustered in *P* dimensional space, where *P* equals the number of principal components explaining >90% of the variance in the dataset. We then performed multiple iterations of *k*-means clustering, varying the value of *k* from 2 to 10 using the *kmeans* function in MATLAB. We calculated the sum of squared distances for each point to its assigned centroid, inspected silhouette plots utilizing the *silhouette* function in MATLAB, and determine the total and mean number of negative silhouette coefficients across all values of *k* to identify the best value. Utilizing the same dataset, we independently performed hierarchical clustering using the ‘ward’ method based on Euclidean distances via the *pdist* and *linkage* functions in MATLAB.

## RESULTS

### Monocularly and binocularly innervated neurons are found throughout the mouse SC

To label neurons in the SC receiving contra-and/or ipsi-RGCs, we utilized an intersectional, transsynaptic tracing strategy (Fig. 1A). First, we leveraged the ability of serotype 1 adeno-associated virus (AAV1) to be transferred trans-synaptically in the anterograde direction by intraocular injection of AAV1-Cre and AAV1-FlpO to express these recombinases in retinorecipient SC neurons^26^. Next, we stereotactically injected independent Cre-and FlpO-dependent AAVs encoding fluorescent reporters with distinct emission spectra bilaterally into the SC. Thus, in each SC hemisphere neurons innervated monocularly by contra-or ipsi-RGCs (SC-C, SC-I) would express distinct reporters, while those binocularly innervated would express both (SC-B). Since the mouse SC is innervated by 20-fold more contra-RGCs than ipsi-RGCs^27^, identification of which hemisphere was contralateral and ipsilateral to a respective ocular injection site was easily determined (Fig. 1B). In many cases, neurons of more than one innervation pattern could be identified in the same image volume (Fig. 1C & D) and SC-C, SC-I, and SC-B cell bodies were readily isolated for determining relative proportions (Fig. 1E & F). Importantly, we randomized which eye received a given virus, as well as which fluorescent reporter was Cre-or FlpO-dependent, and obtained similar results, regardless of the combination of viruses used (Fig. S1).

**Figure 1.**
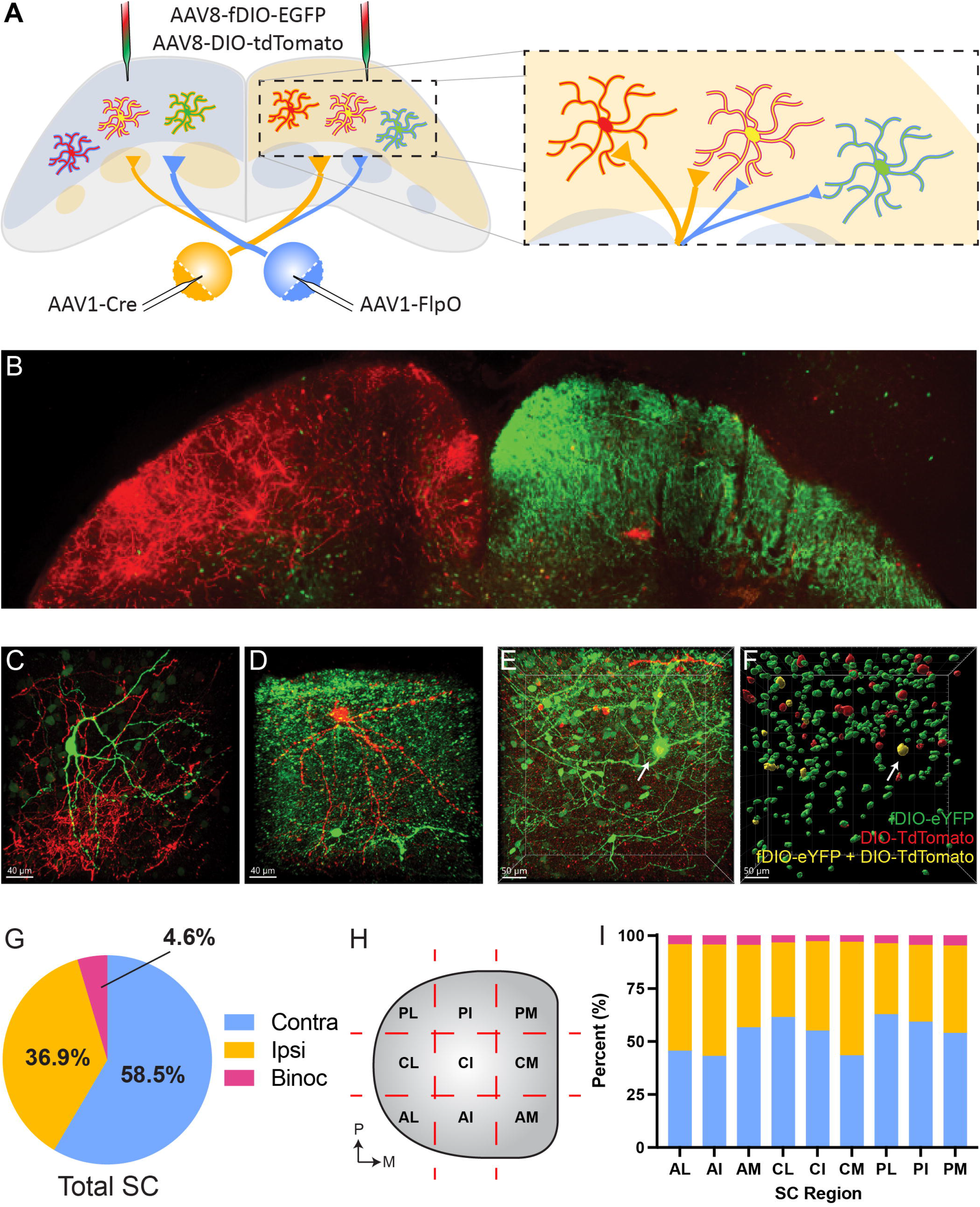
– Monocularly and binocularly innervated neurons are present throughout the SC. A) Schematic of trans-synaptic viral tracing strategy, where monocularly innervated neurons will be labeled with a single fluorophore and binocularly innervated neurons labeled with two fluorophores. B) Widefield image of a coronal section through the SC following trans-synaptic viral tracing. C & D) Representative high-magnification images of SC regions containing neurons innervated by each eye. E & F) Representative volume of the SC containing monocularly and binocularly innervated neurons (E) and automated labeling of each population for quantification. G) Pie chart quantifying the relative proportion of neurons receiving monocular input from contralateral retinal ganglion cells (RGCs) (*blue*), ipsilateral RGCs (*yellow*), or both (*red*). H) Schematic of SC divided into 9 subregions for topographic analysis. I) Bar graph quantifying the relative proportions of neurons receiving input from contralateral RGCs (*blue*), ipsilateral RGCs (*yellow*), or both (*red*). *P, posterior; M, medial; AL, antero-lateral; AI, antero-intermediate; AM, antero-medial; CL, centro-lateral; CI, centro-intermediate; CM, centro-medial; PL, postero-lateral; PI, postero-intermediate; PM, postero-medial*

Having established our labeling paradigm, we first asked what proportion of retinorecipient neurons were innervated by contra-and/or ipsi-RGCs. Consistent with denser contra-RGC input, we found that the majority of labeled neurons were SC-C (58.6 ± 12.3%, N = 22,731 neurons, 8 mice) (Fig. 1G). Surprisingly, even though only ∼5% of RGCs project ipsilaterally^27^, we found that 36.9 ± 11.4% retinorecipient neurons were SC-I and another 4.6 ± 2.2% were SC-B (Fig. 1G). Since ipsi-RGC input is restricted to the antero-medial crescent of the SC, we next asked if the proportions of innervation types varied with topographic position. To do so, we divided the SC into 9 segments spanning the anterior-posterior and medial-lateral axes (Fig. 1H). Strikingly, we found that the relative proportions of SC-C, SC-I, and SC-B neurons were consistent across the entire SC (Fig. 1I). Together, these data demonstrate that distinct patterns of retinal innervation exist in the SC, that the number of SC neurons receiving contra-and ipsi-RGC innervation is not proportional to the relative inputs, and that the relative proportions of neurons receiving each innervation pattern is independent of topographic position.

### SC neurons receiving different innervation patterns have similar morphological characteristics

Since we found relatively similar proportions of SC-C, SC-I and SC-B neurons across the entire SC, we sought to determine if there were morphological differences between these groups. We reconstructed neurons innervated by contra-and/or ipsi-RGCs across the entire SC and quantified an array of morphological parameters. We first examined the size of each population by determining the soma volume, total dendritic length, and convex hull volume. Interestingly, we found that the soma volumes SC-I neurons were significantly smaller than both SC-C and SC-B neurons (P < 0.0001, Kruskal-Wallis and Dunn’s multiple comparisons tests) (Fig. 2A). However, we found no differences in total dendrite length nor convex hull volume between groups (Fig. 2B & C). We next examined dendritic branching patterns of neurons in each group. While we found no differences between groups in the absolute numbers of branches in primary through quinary order, we did find a small but significant decrease in DBOI in SC-B neurons compared to SC-I neurons (P = 0.0409) (Fig. 2D & E). Interestingly, we found that the average primary dendrite diameter of SC-I neurons was decreased compared to SC-C neurons (P = 0.0007) (Fig. 2F).

**Figure 2.**
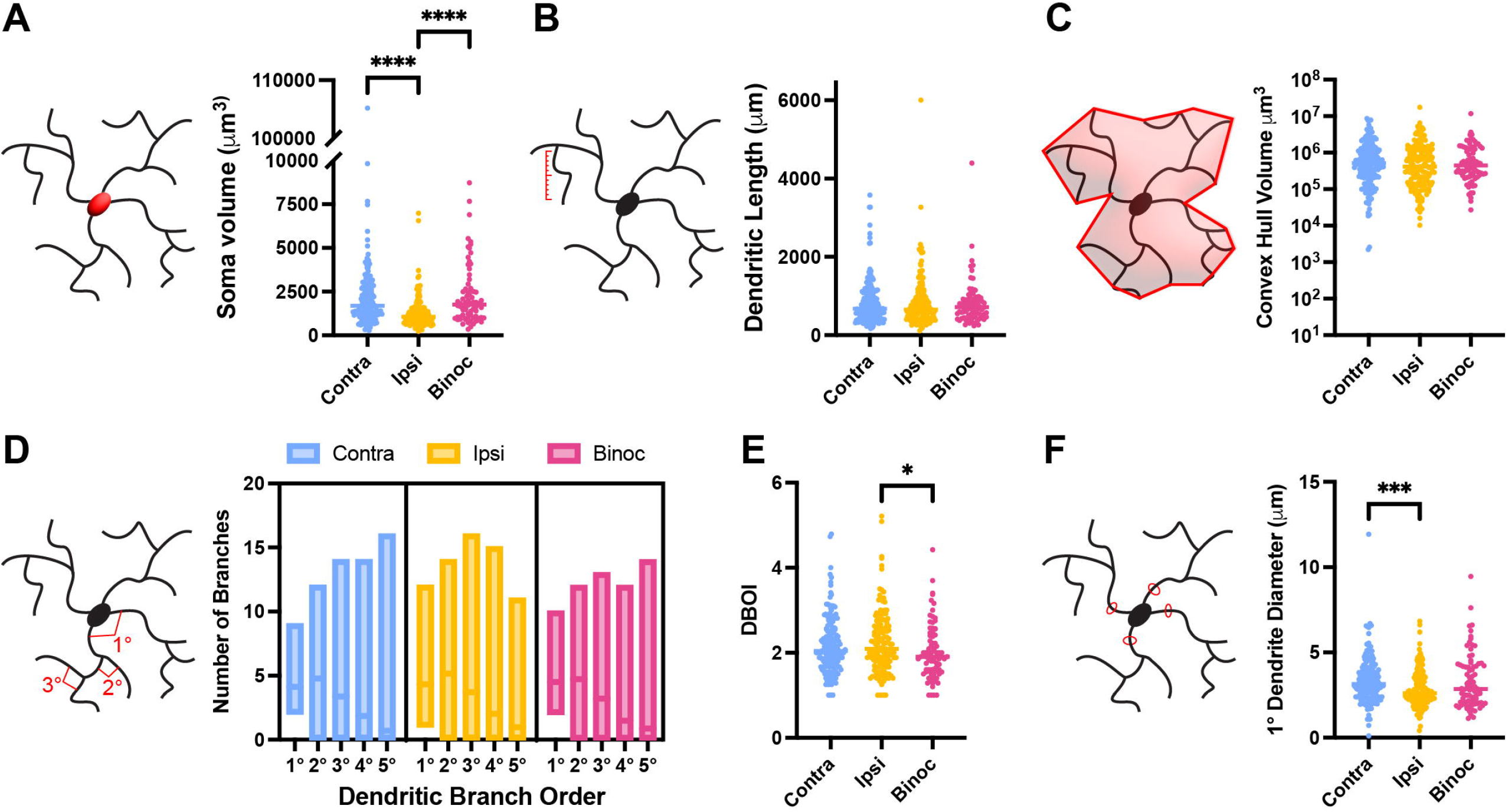
– Basic morphometric features are similar in monocularly and binocularly innervated SC neurons. A-F) Schematics and quantifications of soma volume (A), total dendritic length (B), convex hull volume (C), number of primary through quinary dendritic branches (D), dendritic branch order index (DBOI) (E), and average primary dendrite diameter (F) in SC neurons innervated by contralateral retinal ganglion cells (RGCs) (*blue*), ipsilateral RGCs (*yellow*), or both (*red*). *\*, P < 0.05; ***, P < 0.001; ****, P < 0.0001, Kruskal-Wallis test and Dunn’s multiple comparisons test*

To further explore potential morphological distinctions between differentially innervated neurons in the SC, we performed a Sholl analysis^28^ (Fig. 3A & B). We found no difference in the distribution of dendrite intersections as a function of Sholl radii, the maximum number of intersections, or the radius at which this occurred (Fig. 3B-D). Intriguingly, we did observe a slight decrease in SRI, which is the ratio of the maximum number of Sholl intersections divided by the number of primary dendrites, for SC-B neurons in comparison to SC-I neurons (P = 0.0440) (Fig. 3E). To further quantitatively assess morphology, we grouped Sholl intersections into those occurring in four equal quadrants dorsal, ventral, lateral, and medial relative to the cell soma. This allowed us to determine the height and width of each neuron’s dendritic arbors, as well as the AOI to describe the overall dendrite shape; however, we found no differences between groups for any of these measures. Taken together, these data suggest that, aside from decreased soma volume in SC-I neurons and incompletely penetrant differences in primary dendrite diameter, DBOI, and SRI, the morphologies of SC-C, SC-I, and SC-B neurons are largely similar.

**Figure 3.**
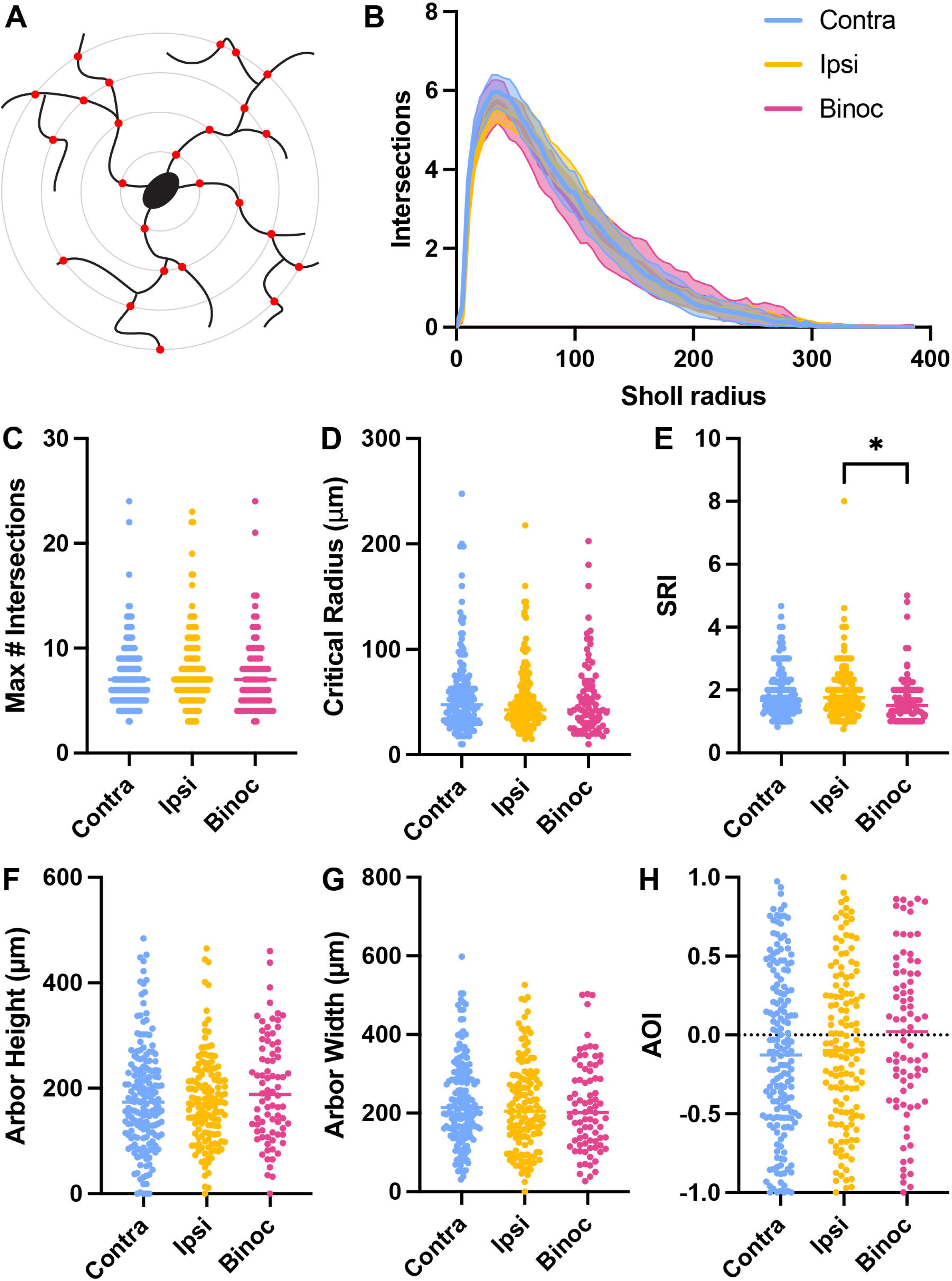
–Sholl-derived morphometric features are similar in monocularly and binocularly innervated SC neurons. A) Schematic of Sholl intersection analysis. B) Quantification of the number of intersections at 5 micron intervals in SC neurons innervated by contralateral retinal ganglion cells (RGCs) (*blue*), ipsilateral RGCs (*yellow*), or both (*red*). C-H) Quantifications of the maximum number of intersections (C), critical radius at which the maximum occurred (D), Sholl ramification index (SRI) (E), dendritic arbor height (F), dendritic arbor width (G), and arbor orientation index (AOI) (H) in SC neurons innervated by contralateral RGCs (*blue*), ipsilateral RGCs (*yellow*), or both (*red*). *\*, P < 0.05, Kruskal-Wallis test and Dunn’s multiple comparisons test*

### Minimal region-specific differences in SC neuron morphology

Since the SC is topographically organized, with central visual field represented mediolaterally, we compared the morphology of SC neurons across topographic subregions of the SC. Interestingly, we did find main effects of region on multiple parameters (DBOI, P = 0.001; maximum Sholl intersections, P = 0.0104; SRI, P = 0.0134; arbor width, P < 0.0001; AOI, P < 0.0001; 2-way ANOVA). DBOI was decreased for neurons in AM compared to those in centrolateral (CL) and posterointermediate (PI) regions of SC (Fig. S2). The maximum number of Sholl intersections was increased for neurons in centrolateral (CL) compared to those in AL and anteromedial (AM) regions of SC, while SRI was increased for neurons in CL compared to those in AM (Fig. S3). In addition, arbor width was decreased, while AOI was increased, in neurons in CM and PM regions compared to those in AL, AI, PL, and PI regions (Fig. S3). These data raised the possibility that there may be regional differences in morphology between neurons receiving different innervation patterns.

We compared the previously described morphological characteristics of SC-C, SC-I, and SC-B neurons across nine subsections of the SC across the medio-lateral (M-L) and antero-posterior (A-P) axes (Fig. 4). Consistent with our previous analysis, we found main effects of innervation type on average primary dendrite diameter (P = 0.009, 2-way ANOVA), with SC-I neurons having smaller diameters (P = 0.0174, SC-I vs. SC-C; P = 0.0379, SC-I vs. SC-B; Tukey’s multiple comparisons test), and DBOI (P = 0.0356), with SC-B neurons having smaller values than SC-I neurons (P = 0.028). However, neither of these effects were observed within a given subregion, instead SC-B neurons in anterolateral (AL) SC had larger primary dendrite diameters than SC-C and SC-I neurons (Fig. 4E & F). In contrast to our previous analysis, no innervation type effects were observed for soma volume (P = 0.1149) or SRI (P = 0.1192), suggesting the differences previously observed are not as robust. Indeed, we found within region effects of innervation type in only a few cases. We observed increased soma volume in SC-C neurons in AM, fewer primary branches in SC-I neurons in posteromedial (PM), and increased primary dendrite diameter in SC-B neurons in AL regions of SC (Fig. 4A, D, & F). In addition, we found a greater number of maximum Sholl intersections in SC-I neurons in anterointermediate (AI), increased arbor width for SC-B neurons in AI and SC-C neurons in centromedial (CM), and increased AOI for SC-B neurons in posterolateral (PL) regions of the SC (Fig. 4G, K & L). Furthermore, no interactions between innervation type and region were observed for any metric measured. Together, these data suggest that some regional differences in morphology exist within the SC, but that within a given region the morphologies of SC-C, SC-I, and SC-B neurons are largely similar.

**Figure 4.**
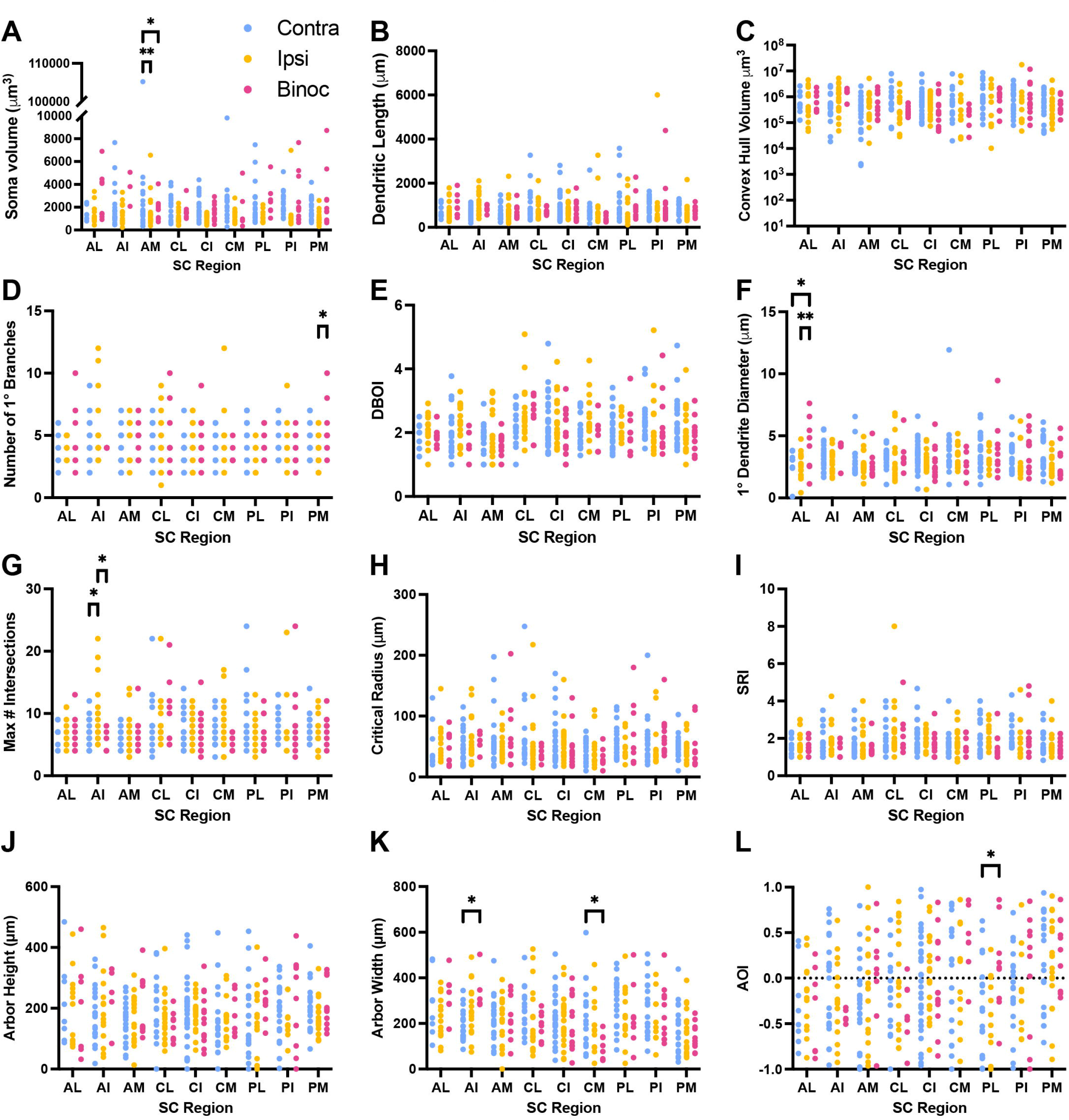
– Morphometric features of monocularly and binocularly innervated neurons are similar across topographically-distinct regions of the SC. A-L) Quantifications of soma volume (A), total dendritic length (B), convex hull volume (C), number of primary through quinary dendritic branches (D), dendritic branch order index (DBOI) (E), average primary dendrite diameter (F), the maximum number of intersections (G), critical radius (H), dendritic arbor height (I), dendritic arbor width (J), Sholl ramification index (SRI) (K), and arbor orientation index (AOI) (L) in SC neurons innervated by contralateral retinal ganglion cells (RGCs) (*blue*), ipsilateral RGCs (*yellow*), or both (*red*) in different subregions of the SC. *AL, antero-lateral; AI, antero-intermediate; AM, antero-medial; CL, centro-lateral; CI, centro-intermediate; CM, centro-medial; PL, postero-lateral; PI, postero-intermediate; PM, postero-medial; *, P < 0.05; **, P < 0.01, 2-way ANOVA and Tukey’s multiple comparisons test*

### Identification of five morphologically distinct subtypes in the SC

Though we found minimal differences in the morphometric features due to innervation pattern or topographic location, this could be driven by the complement of morphological subtypes comprising these populations. Previous studies identified 4-6 different morphological subtypes in the SC^15–17^, and we sought to determine if a similar classification could be derived from our dataset. To do so, we utilized 10 morphometric features and performed principal components analysis to reduce dimensionality. We subsequently utilized *k*-means clustering to identify five morphologically distinct subtypes in our dataset (N = 388 neurons from 8 mice) (Fig. 5A). To determine the optimal value for *k*, we performed multiple iterations of clustering for values ranging from two to ten. By plotting the sum of the squared distances from each point to its assigned centroid as a function of the value of *k*, we observed substantial reductions up to a value of five, with smaller reductions thereafter (Fig. 5B). Additionally, we noted decreases in the total and average number of negative silhouette coefficients when *k* increased from four to five (Fig. 5C), confirming that five clusters describe the data more accurately. We next determined the relative proportions of each cluster, which we named in descending order of prevalence (Fig. 5D). Notably, three clusters accounted for nearly 82% of the population (Cluster 1, 29.12%; Cluster 2, 28.87%; Cluster 3, 23.97%), while Clusters 4 and 5 accounted for 11.86% and 6.19% of the population, respectively. Visual inspection of reconstructed neurons from each cluster confirmed their distinctness and aligned with previously published analyses in the SC (Fig. 5E). Cluster 1 appeared similar to small stellate cells (Fig. 5E, *blue*), Cluster 2 resembled narrow field cells (Fig. 5E, *red*), Cluster 3 were akin to larger stellate cells (Fig. 5E, *green*), Cluster 4 approximated horizontal cells (Fig. 5E, *purple*), and Cluster 5 appeared similar to widefield cells (Fig. 5E, *orange*).

**Figure 5.**
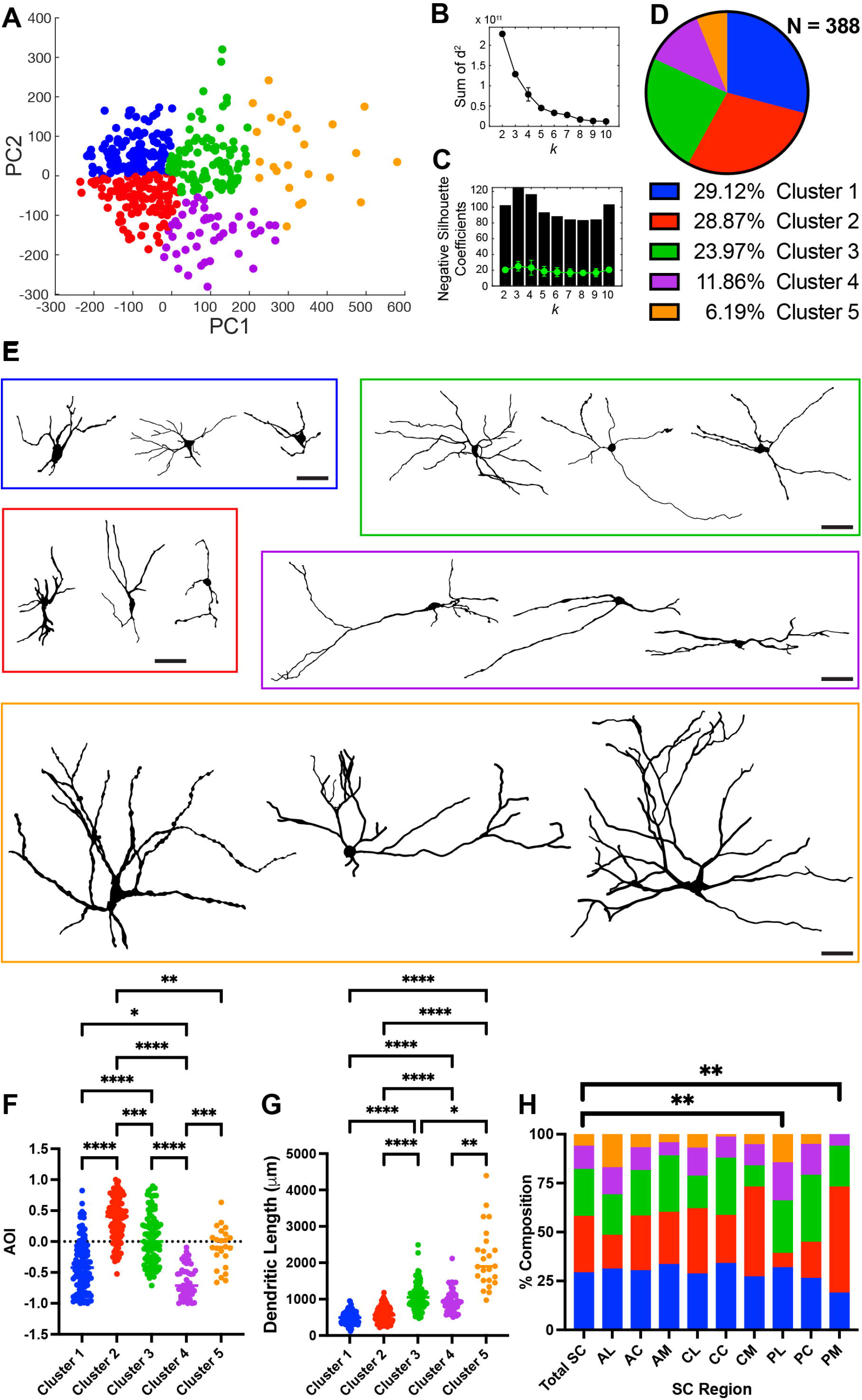
– Different morphological subtypes are evenly distributed throughout the SC. A) Scatter plot of first and second principal components (PC) based on 10 morphometric features. Different colors label distinct clusters identified via *k*-means clustering. B & C) Quantification of the sum of squared distances to centroids (B), as well as total and average number of negative silhouette coefficients (C) suggest 5 is the optimal number of clusters following iterative *k-*means clustering using values 2-10 for *k*. D) Pie chart quantifying the relative proportions of clusters. E) Representative reconstructions of neurons belonging to each cluster. *Bar, 50 μm* F & G) Quantification of arbor orientation index (F) and total dendritic length (G) in identified clusters. *\*, P < 0.05; **, P < 0.01; ***, P < 0.001; ****, P < 0.0001, Kruskal-Wallis test and Dunn’s multiple comparisons test.* H) Bar graph quantifying the relative proportions of each cluster throughout the entire SC and in each subregion. *AL, antero-lateral; AI, antero-intermediate; AM, antero-medial; CL, centro-lateral; CI, centro-intermediate; CM, centro-medial; PL, postero-lateral; PI, postero-intermediate; PM, postero-medial; **, P < 0.01, Chi-square test*

To assess the validity of our cluster segregation, we determine if there were differences in morphometric features between groups. As expected, we found significant differences between clusters for all features included in our clustering algorithm (data not shown). In addition, we found significant differences between clusters for morphometric features not included in the clustering algorithm, such as soma volume, total dendritic length, and convex hull volume (P < 0.0001, Kuskal-Wallis test) (Fig. 5F & G and data not shown). We also performed independent hierarchical clustering, which similarly identified five well separated clusters and a substantial degree of overlap with *k*-means based clusters (Fig. S4).

Given our previous identification of regional differences in morphology (Figs. S2 & S3), we next asked if there were biases in the distribution of these morphological subtypes across topographic subregions of the SC. In comparison to the whole SC, the relative proportions of each morphological subtype were similar in all subregions except the PL and PM SC (P = 0.0092, PL vs. total SC; P = 0.0021, PM vs. total SC; Chi-square test) (Fig. 5H). In PL, there was a paucity of Cluster 2, while in PM there was an over-representation of Cluster 2 and a lack of Cluster 5. Taken together, these data suggest that there are multiple morphologically distinct subtypes in the SC, which are distributed largely evenly across topographic subregions.

### Monocularly and binocularly innervated neurons are comprised of the full diversity of morphologic subtypes

We next asked if SC-C, SC-I, and SC-B neurons were comprised of different proportions of morphologically distinct subtypes. Visual inspection of reconstructed neurons of each innervation pattern revealed a heterogeneous array of morphologies (Fig. 6A-C). Indeed, quantification revealed that all identified subtypes could be found amongst SC-C, SC-I, and SC-B neurons (Fig. 6A-D). Furthermore, the relative proportions of subtypes were strikingly similar across innervation pattern and did not differ from the distribution across the entire SC (Fig. 6D). We next asked if there were a bias in the innervation pattern composition of each morphologic subtype. Consistent with the even distribution of morphologies across innervation types, we observed similar compositions of innervation pattern across morphologic subtypes (Fig. 6E).

**Figure 6.**
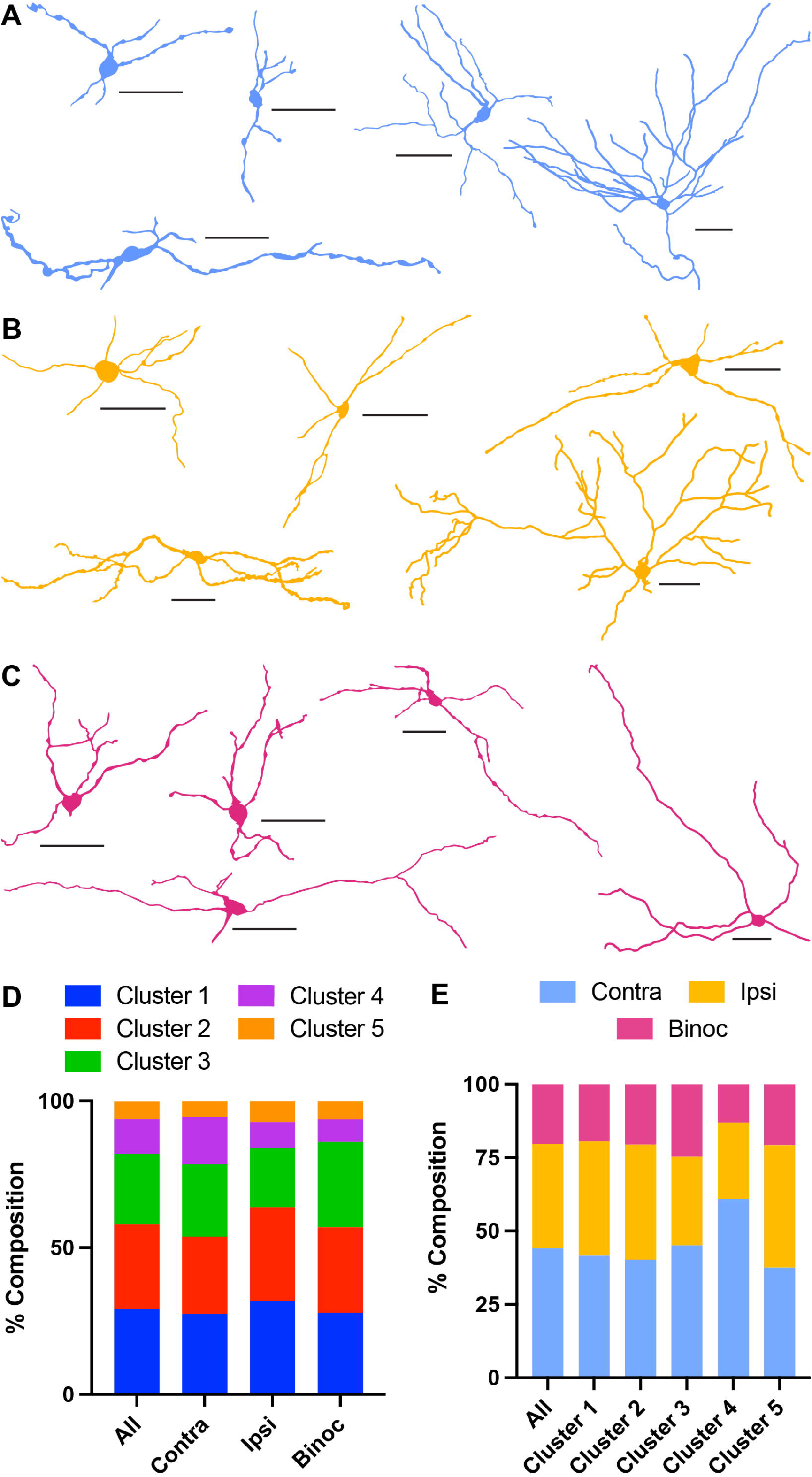
– Monocularly and binocularly innervated SC neurons are comprised of all morphological subtypes. A-C) Representative examples of reconstructed SC neurons innervated by contralateral retinal ganglion cells (RGCs) (A), ipsilateral RGCs (B), or both (C). D) Bar graph quantifying the relative proportions of each morphological subtype across all reconstructed SC neurons and each type of innervation pattern. E) Bar graph quantifying the relative proportions of SC neurons innervated by contralateral RGCs, ipsilateral RGCs, or both across all reconstructed neurons and each morphological subtype.

To further interrogate any potential effects of retinal innervation pattern on morphology, we asked if there were any differences in morphometric features between morphologic subtypes. Somewhat surprisingly, we found innervation pattern-driven differences in several features (Fig. S5). Specifically, we found a main effect of innervation type for soma volume (P < 0.0001, 2-way ANOVA), convex hull volume (P = 0.0299), SRI (P = 0.0453), and average primary dendrite diameter (P = 0.0014). We also observed an interaction between innervation pattern and morphologic subtype for dendritic length (P = 0.0277), the maximum number of Sholl intersections (P = 0.0397), critical radius (P = 0.0442), average primary dendrite diameter (P = 0.0466), and arbor width (P = 0.0310). However, there did not appear to be a consistent morphologic subtype that exhibited innervation-driven differences. Interestingly, *post hoc* multiple comparisons tests revealed that most differences were between SC-I neurons and either SC-C or SC-B neurons, and we found only one difference between SC-C and SC-B neurons. Taken together, these data suggest that there may be differences in morphometric features within a morphologic subtype due to innervation pattern but that SC-C, SC-I, and SC-B neurons are comprised of similar morphological subtypes.

### Overlapping expression of molecular markers across SC-C, SC-I, and SC-B neurons

In addition to morphologic features, SC subtypes can be identified based on gene expression patterns. To determine if there may be molecular differences between SC neurons receiving different eye-specific innervation patterns, we co-labeled for proteins known to be expressed in subsets of SC neurons. We found similar proportions of SC-C, SC-I, and SC-B neurons expressed the markers Tal1, parvalbumin, calbindin, and Meis2 (N > 1000 cells per animal from 7 animals) (Fig. 7). Together with our previous findings of minimal morphologic differences between and overlapping compositions of morphologic subtypes in SC-C, SC-I, and SC-B neurons, these data suggest that similar populations of SC neurons receive all patterns of eye-specific input.

**Figure 7.**
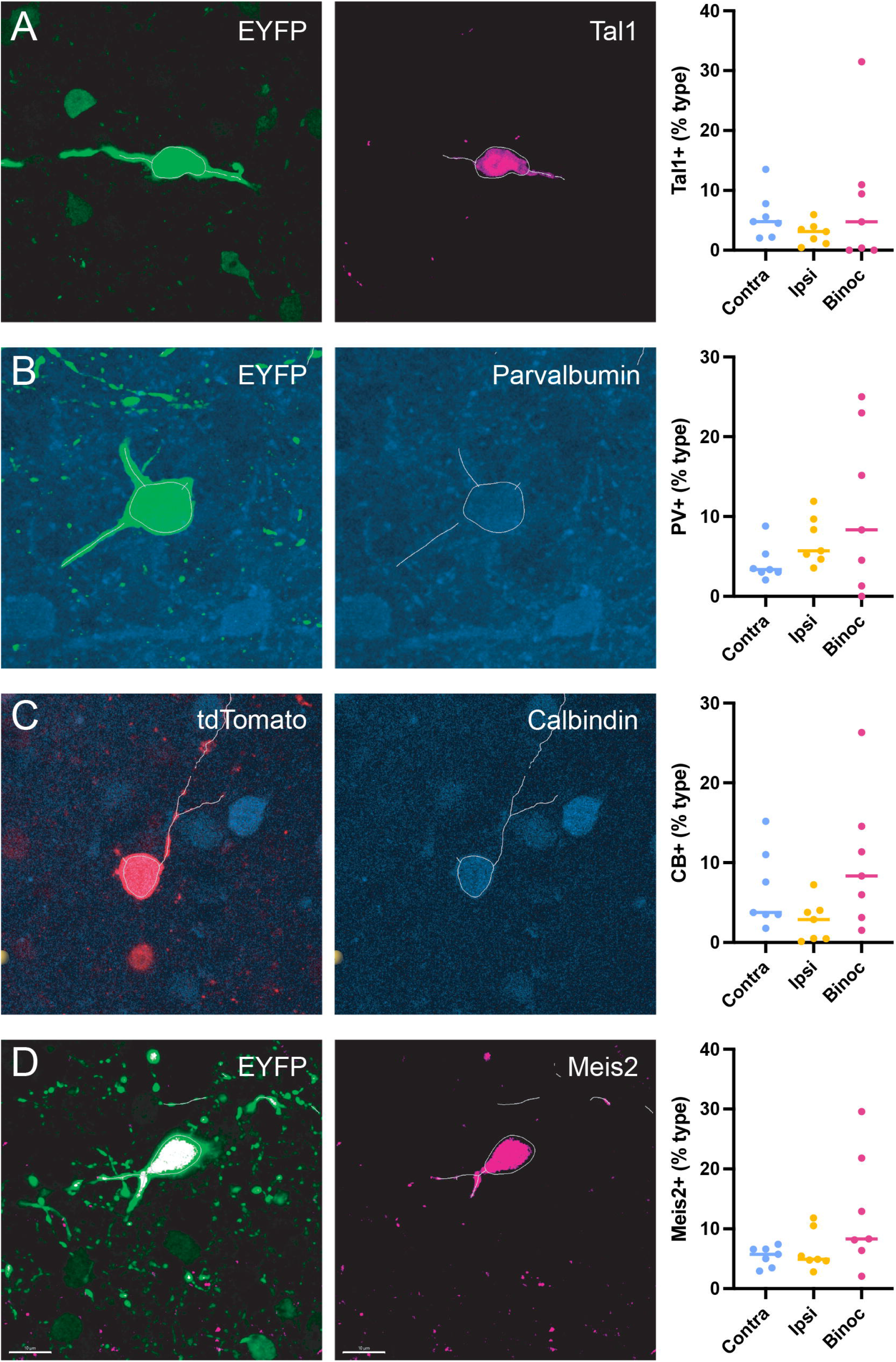
– Similar expression of candidate molecular markers in monocularly and binocularly innervated neurons. A-D) Photomicrographs of neurons expressing a fluorescent marker induced by trans-synaptic tracing (*left*) and candidate molecular markers (*middle*) Tal1 (A), parvalbumin (B), calbindin (C), or Meis2 (D), as well as quantification of the percent of trans-synaptically labeled neurons expressing the marker (*right*).

## DISCUSSION

Recent studies suggest that binocular interactions in the mouse SC are more prevalent than previously thought and that binocular vision is critical for ethologically relevant behaviors mediated by the SC. The SC receives direct innervation from both contra-and ipsi-RGCs, which could relay eye-specific information in three ways: along distinct parallel channels to specific neuronal subtypes, along shared pathways to all neuronal subtypes, or a combination thereof. Here, we utilized an intersectional, trans-synaptic tracing strategy to label neurons receiving different patterns of eye-specific innervation and find that each targets overlapping subsets of SC neurons defined by morphology and candidate marker expression. Interestingly, we also find that eye-specific innervation patterns are represented across the entire SC. Together, these data suggest that eye-specific information is relayed in a distributed manner to the SC and multiplexed with feature-specific information to inform behavioral outputs.

### Distribution of eye-specific information across overlapping subtypes

In the mouse SC, RGC inputs are organized based on eye of origin, location within the retina, and functional specification. Whether and how these organizations interact shape the processing capabilities of the SC^29^. On one hand, highly specialized channels of information could enable rapid visuomotor transformations. In support of this possibility, previous studies suggest that orientation preference could be represented differentially in topographically segregated regions of the SC^30,31^. Alternatively, broad distribution of tuning capabilities would allow for complete feature space to be encoded universally. In support of this, Off-α RGCs capable of triggering an escape behavior in response to a looming stimulus are distributed widely across the retina^32^. Given the recent characterization of binocular interactions in the mouse SC, we know little about how eye-specific information is relayed to inform visual processing.

The SC plays a critical role in ethologically relevant behaviors, including prey capture^5,6^ and visual threat avoidance^7^, both of which are dependent on binocular vision^2,3^. Given the importance of these behaviors to survival, dedicated retinocollicular subcircuits relaying stimulus and eye-specific information could be advantageous. Alternatively, the addition of eye-specific context broadly across circuits utilized for visual processing in the SC could increase the feature space encoded. We find that the latter strategy is utilized in the mouse SC, as overlapping morphologic and molecular subtypes are found in similar proportions across eye-specific innervation patterns. However, it should be noted that SC-B neurons represent only ∼5% of the population, while ∼60% of SC neurons exhibit binocular modulation^12–14^. Thus, alternate paths of photic relay from the ipsilateral eye likely underlie a substantial proportion of binocular processing in the SC^8^. While the findings presented here add to our understanding of how visual information is organized in the SC, future studies utilizing methods to determine the organization of alternate binocular convergence in the SC are needed.

Previous work identified four morphologic subtypes in the SC and suggested a strong relationship between morphology, intrinsic electrophysiological properties, and visual response properties in the SC^17^. However, substantial variability in tuning were present in each morphologically defined type and, as with all interrogations of visual function, the feature space sampled was limited by the range of stimuli shown. Thus, it is likely that SC neurons could be subdivided into a wider range of classes. Indeed, recent analysis using a battery of six different types of visual stimulus uncovered 24 functionally distinct classes in the mouse SC^23^. And, single nucleus RNA sequencing of the superficial SC revealed 28 transcriptomically distinct classes^20^. Remarkably, this study also combined calcium imaging with *post hoc* visualization of molecular markers to correlate functional and transcriptional identities, moving the field substantially closer to establishing a unified understanding of neuronal diversity in the SC. Our findings that eye-specific information is equally distributed across all morphologic types raise the possibility that the ocularity of responses in the SC further diversify the information coded by this critical visual nucleus.

### Widespread innervation of SC neurons by ipsi-RGCs

A surprising finding from these experiments was the disproportionate representation of ipsi-RGC input in the SC. Our data suggest that ∼40% of SC neurons are directly innervated by ipsi-RGCs, despite them representing only 5% of the total RGC population. Furthermore, we find that SC-I neurons are encountered in similar proportions throughout the SC, even though ipsi-RGC terminals are predominantly localized to the anterior and medial borders of the SC. At first blush, these data seem to upend the well-established topographic order of retinocollicular projections, as ipsi-RGCs are thought to be localized to the ventro-temporal crescent of the retina^2,27^. However, recent work utilizing a genetic method to mark ipsi-RGCs (*Sert-Cre*) revealed that labeled cells occupied ∼30% of the retinal surface area in adults and that a smattering of cells could be found outside the ventro-temporal crescent in adults^33^. Additionally, *in vivo* electrophysiological approaches also reveal responses to ipsilateral eye stimulation across the SC^14^, raising the possibility that ipsi-RGCs could influence activity despite sparse innervation. To determine the relative influence of contra-and ipsi-RGCs across the SC, *in vitro* electrophysiological and optogenetic approaches are needed^34^.

Another possible explanation for the disproportionate representation of ipsi-RGC input in could be the tracing strategy utilized. Our approach leverages trans-synaptic tracing, meaning that neurons receiving only minimal innervation would be labeled as strongly as those receiving dense innervation. If this were the case, we would expect that the dendritic trees of SC-I neurons tend to be larger and span outside of the local topography. However, we did not observe increases in the total dendritic length, convex hull volume, or arbor width of SC-I neurons (Figs. 2 & 3). Furthermore, we did not observe differences in the proportions of morphologic subtypes, which might explain topographically inappropriate connectivity. A final possibility is that the widespread prevalence of SC-I neurons is an artifact of development. We do not think this is the case, as AAV1 injections were made well past the conclusion of RGC topographic refinement in the SC^35^.

### Subtle differences within topographic subregions and morphologic types

While our data suggest that the composition of neurons receiving different patterns of eye-specific input are largely similar, we did uncover some statistically significant differences in a few morphometric features when focusing on topographic subregion or morphologic type. In our interpretation, we did not weigh these differences heavily for two reasons. First, we rarely found instances in which a given morphometric feature was distinct to neurons receiving one innervation pattern. Second, we did not observe consistent differences across regions or subtypes. For instance, soma volume was significantly decreased in SC-I neurons compared to SC-C and SC-B neurons, but this was true only in 1/9 subregions and for only 2/5 morphologic subtypes. However, these region-and subtype-specific differences could play important roles in the visual computations performed by neurons receiving different patterns of eye-specific input. Future studies interrogating the function of SC-C, SC-I, and SC-B neurons in different topographic regions are necessary to resolve this possibility. Related to this, here we only examine morphologic and candidate molecular criteria to define cell types. As mentioned above, utilization of functional and transcriptomic methods to define cell populations reveals more diversity in the SC. Our data do not exclude the possibility that eye-specific information is routed to distinct populations defined by these criteria.

### Identification of five morphologic subtypes

Another surprising finding of this work is the identification of five morphologic subtypes in the SC. Previously, Gale and Murphy sampled a similarly sized population of neurons and uncovered four subtypes: stellate, narrow field, horizontal, and widefield^17^. One potential reason for this could be the differences in metrics chosen for clustering. For example, we chose not to include soma volume, due to the decrease observed in SC-I neurons, or soma distance from the SC surface, because the density of labeling often precluded accurate determination of this metric. Another possibility could be that we under-sampled widefield neurons in our data set, which represented only ∼6% of the population (Fig. 5D). As such, it is possible that more subtle distinctions between the remaining neurons could be identified. Finally, our dataset was substantially larger than that previously analyzed (388 vs. 237 neurons), perhaps allowing additional groups to emerge. Notably, our finding of five subtypes is not out of step with previous work using qualitative metrics to define morphologic types^15^. This study identified six types in the rat SC: stellate, narrow field, horizontal, widefield, marginal, and piriform. Interestingly, Clusters 1, 2, 4, and 5 in our dataset appear to align with stellate, narrow field, horizontal, widefield designations, while Cluster 3 appears similar to either marginal or piriform types.

### Conclusion

Here we utilized a trans-synaptic, intersectional tracing strategy to label neurons in the SC receiving input monocularly from contra-RGCs, monocularly from ipsi-RGCs, and binocularly from both. Interestingly, we find that a substantial portion of retino-recipient neurons are innervated by ipsi-RGCs. Furthermore, we find that the proportions of SC-C, SC-I, and SC-B neurons are similar throughout the SC, suggesting different patterns of eye-specific input are widespread. Additionally, we found few morphometric differences between SC-C, SC-I, and SC-B neurons, suggesting each is comprised by similar morphologic subtypes. Indeed, each of the five morphologic subtypes identified by independent clustering methods are encountered in similar proportions across eye-specific innervation patterns. Finally, we show that the expression of candidate molecular markers does not vary between populations, supporting a model in which eye-specific information is multiplexed on top of other criteria of neuronal identity. Together, these findings add to our understanding of the organization of visual inputs in the SC and set the stage for future investigations interrogating the function of neurons receiving different patterns of eye-specific input.

## Acknowledgements

This work was supported by National Institutes of Health grants R01EY025267 (J.W.T.), R21EY034660 (J.W.T.) and the District of Columbia Intellectual and Developmental Disabilities Research Center (DC-IDDRC) Award P50HD105328, as well as the Waly and Lilly Disney Award for Amblyopia Research from the Research to Prevent Blindness Foundation.

