## Supplemental Figures for "Different patterns of eye-specific input are relayed to overlapping populations in the mouse superior colliculus"

Affiliations: [1]Center for Neuroscience Research, Children's National Hospital, Washington, DC, USA; [2]Neuroscience and Cognitive Science Program, University of Maryland, College Park, MD, USA; [3]Department of Pediatrics, The George Washington University School of Medicine and Health Sciences, Washington, DC, USA; [4]Department of Pharmacology and Physiology, The George Washington University School of Medicine and Health Sciences, Washington, DC, USA

### Figure S1

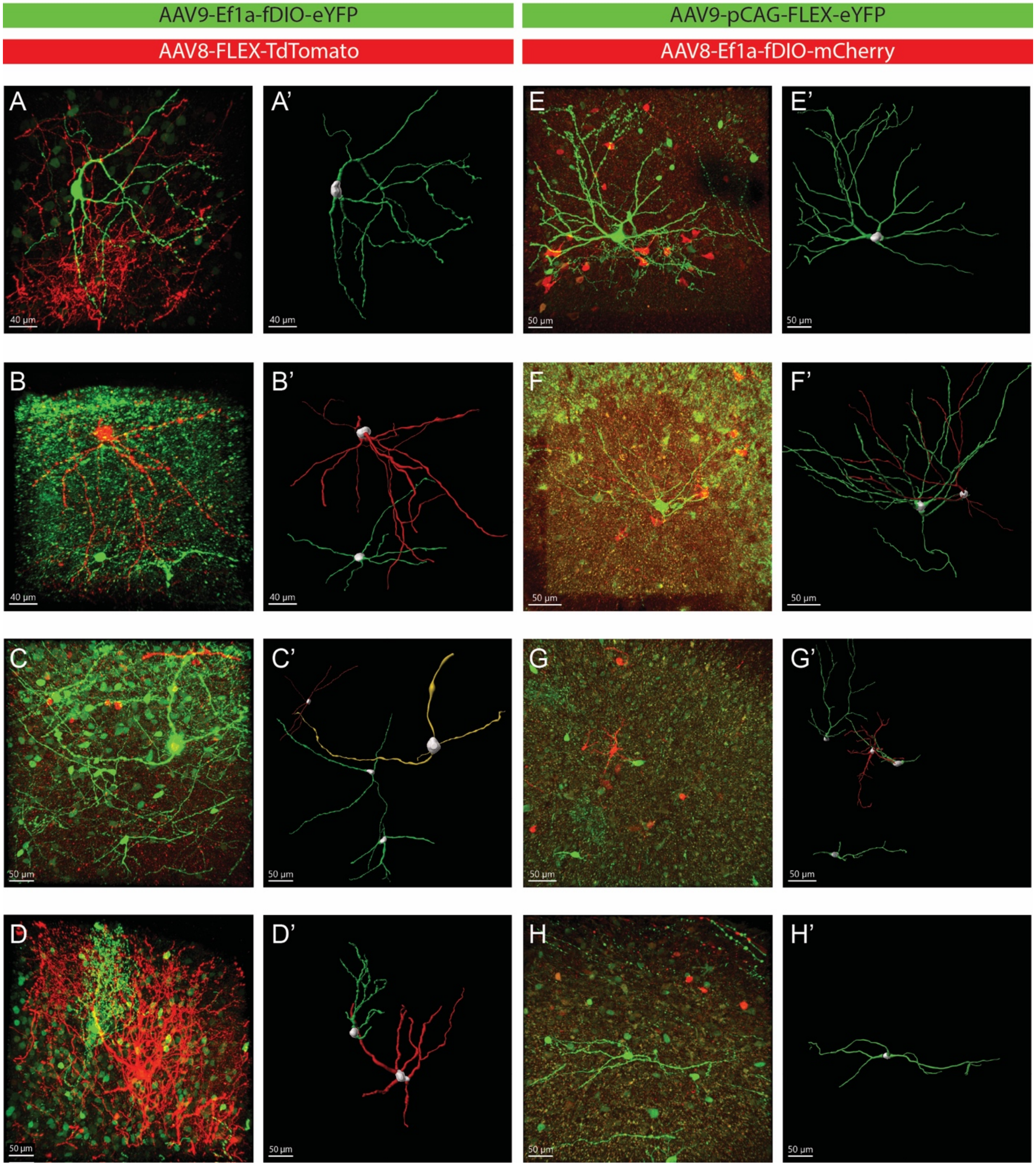

AAV9-Ef1a-fDIO-eYFP
AAV8-FLEX-TdTomato
AAV9-pCAG-FLEX-eYFP
AAV8-Ef1a-fDIO-mCherry
A
A'
E
E'
40 µm
50 µm
B
B'
F
F'
C
C'
G
G'
D
D'
H
H'

**Figure S1 – Trans-synaptic, intersectional labeling of monocularly- and binocularly-innervated SC neurons using alternate combinations of reporter viruses.** A-D) Max intensity projections of confocal image stacks taken at 40X magnification depicting neurons labeled with Cre-dependent red and Flp-dependent green fluorophores. A'-D') Reconstructed neurons from images shown in A-D. E-H) Max intensity projections of confocal image stacks taken at 40X magnification depicting neurons labeled with Flp-dependent red and Cre-dependent green fluorophores. E'-H') Reconstructed neurons from images shown in E-H.

### Figure S2

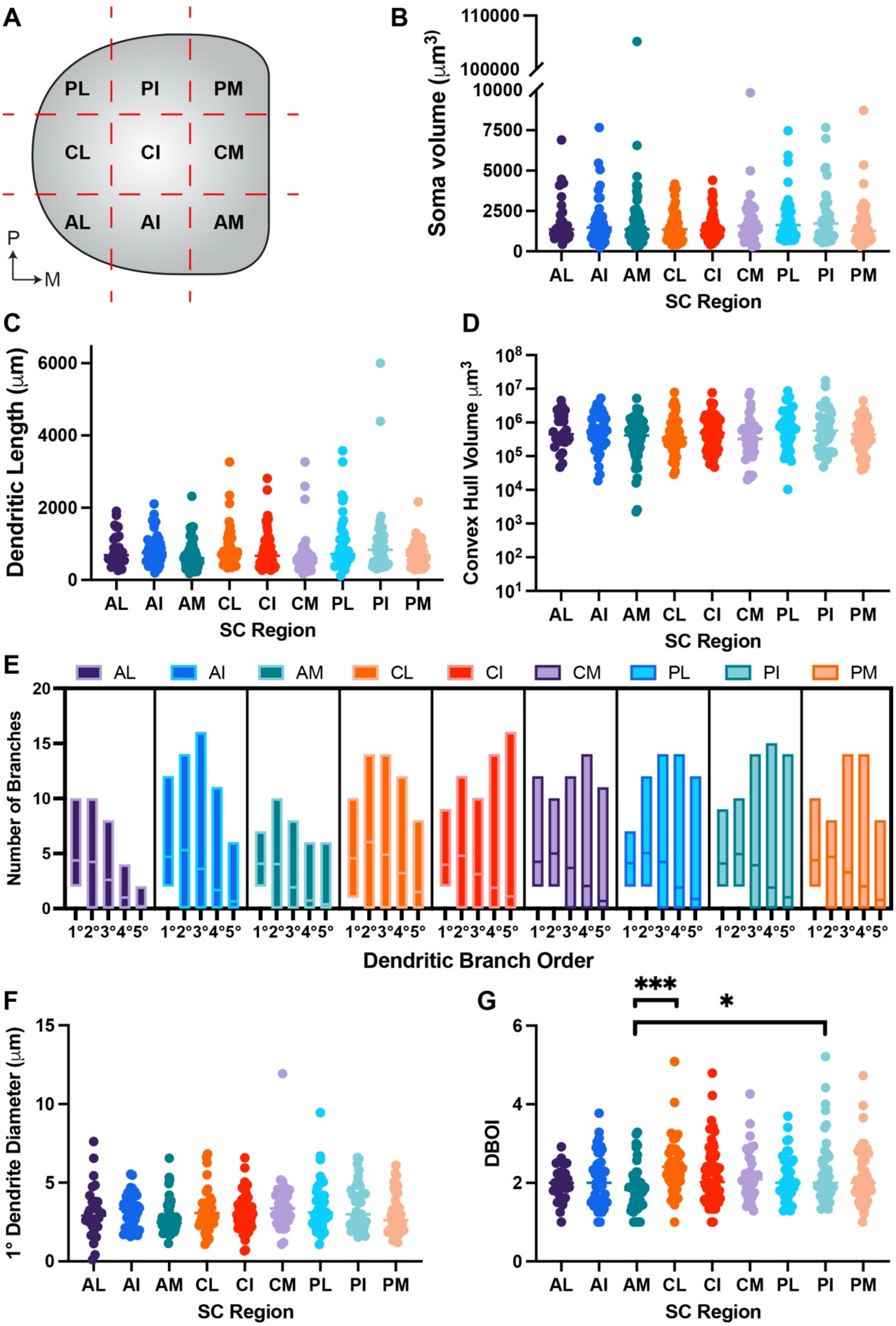

A
PL
PI
PM
CL
CI
CM
AL
AI
AM
P
M
B
Soma volume (μm3)
SC Region
AL AI AM CL CI CM PL PI PM
C
Dendritic Length (μm)
D
Convex Hull Volume μm3
E
Number of Branches
Dendritic Branch Order
1°2°3°4°5°
F
1° Dendrite Diameter (μm)
G
DBOI
***
*

**Figure S2 – Analysis of basic morphological features in different topographic subregions of the SC.** A) Schematic of SC divided into 9 subregions for topographic analysis. B-G) Quantifications of soma volume (B), total dendritic length (C), convex hull volume (D), number of primary through quinary dendritic branches (E), average primary dendrite diameter (F), and dendritic branch order index (DBOI) (E) in SC neurons across different topographic subregions. *P, posterior; M, medial; AL, antero-lateral; AI, antero-intermediate; AM, antero-medial; CL, centro-lateral; CI, centro-intermediate; CM, centro-medial; PL, postero-lateral; PI, postero-intermediate; PM, postero-medial; *, $P < 0.05$; ***, $P < 0.001$, Kruskal-Wallis test and Dunn's multiple comparisons test*

### Figure S3

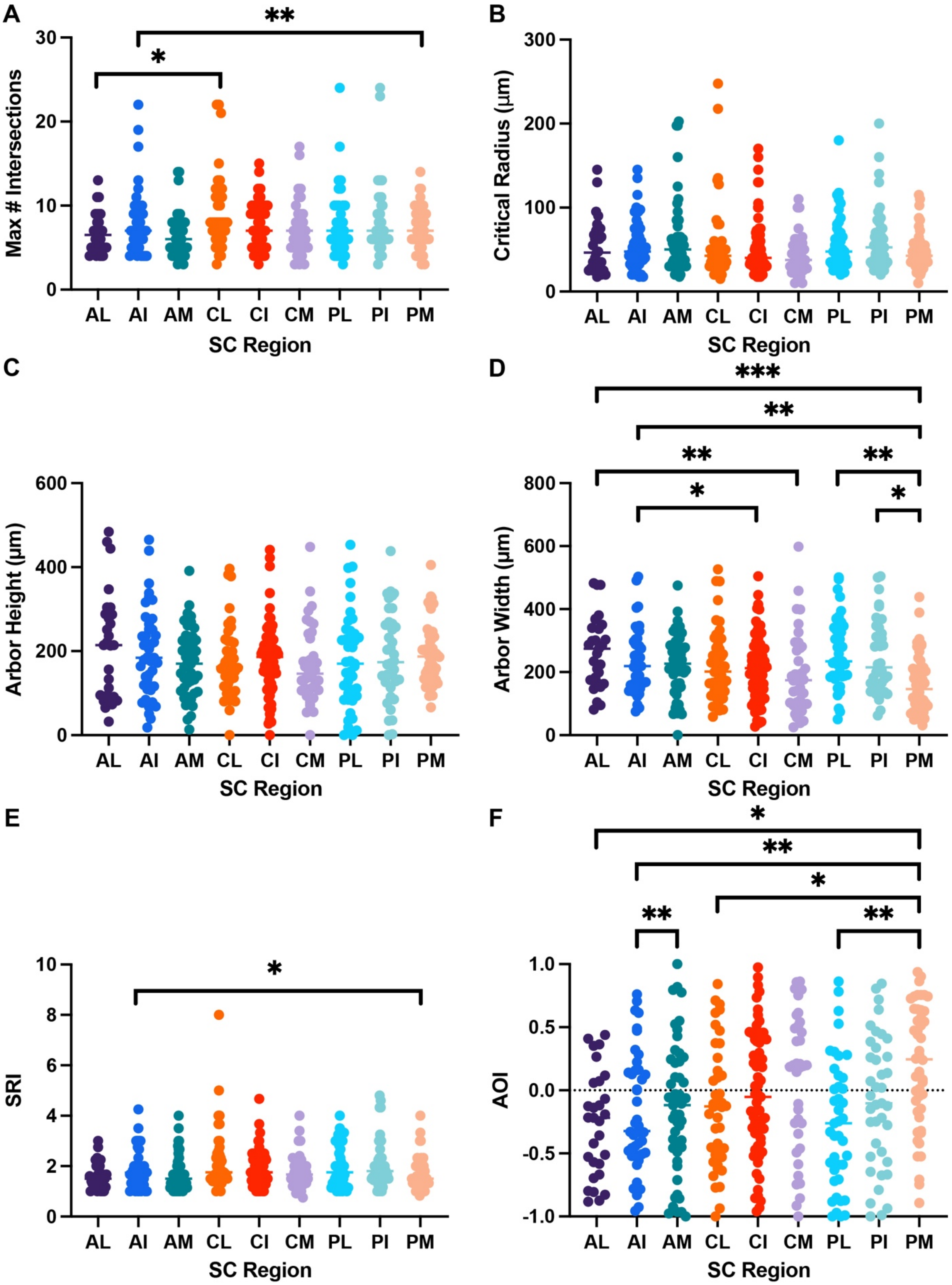
A
B
C
D
E
F
Max # Intersections
Critical Radius (μm)
Arbor Height (μm)
Arbor Width (μm)
SRI
AOI
SC Region
AL
AI
AM
CL
CI
CM
PL
PI
PM

**Figure S3 - Analysis of Sholl-derived morphological features in different topographic subregions of the SC.** A-F) Quantifications of the maximum number of intersections (A), critical radius at which the maximum occurred (B), dendritic arbor height (C), dendritic arbor width (D), Sholl ramification index (SRI) (E), arbor orientation index (AOI) (F) in SC neurons across different topographic subregions. *P, posterior; M, medial; AL, antero-lateral; AI, antero-intermediate; AM, antero-medial; CL, centro-lateral; CI, centro-intermediate; CM, centro-medial; PL, postero-lateral; PI, postero-intermediate; PM, postero-medial;* *, $P < 0.05$; **, $P < 0.01$; ***, $P < 0.001$, *Kruskal-Wallis test and Dunn's multiple comparisons test*

### Figure S4

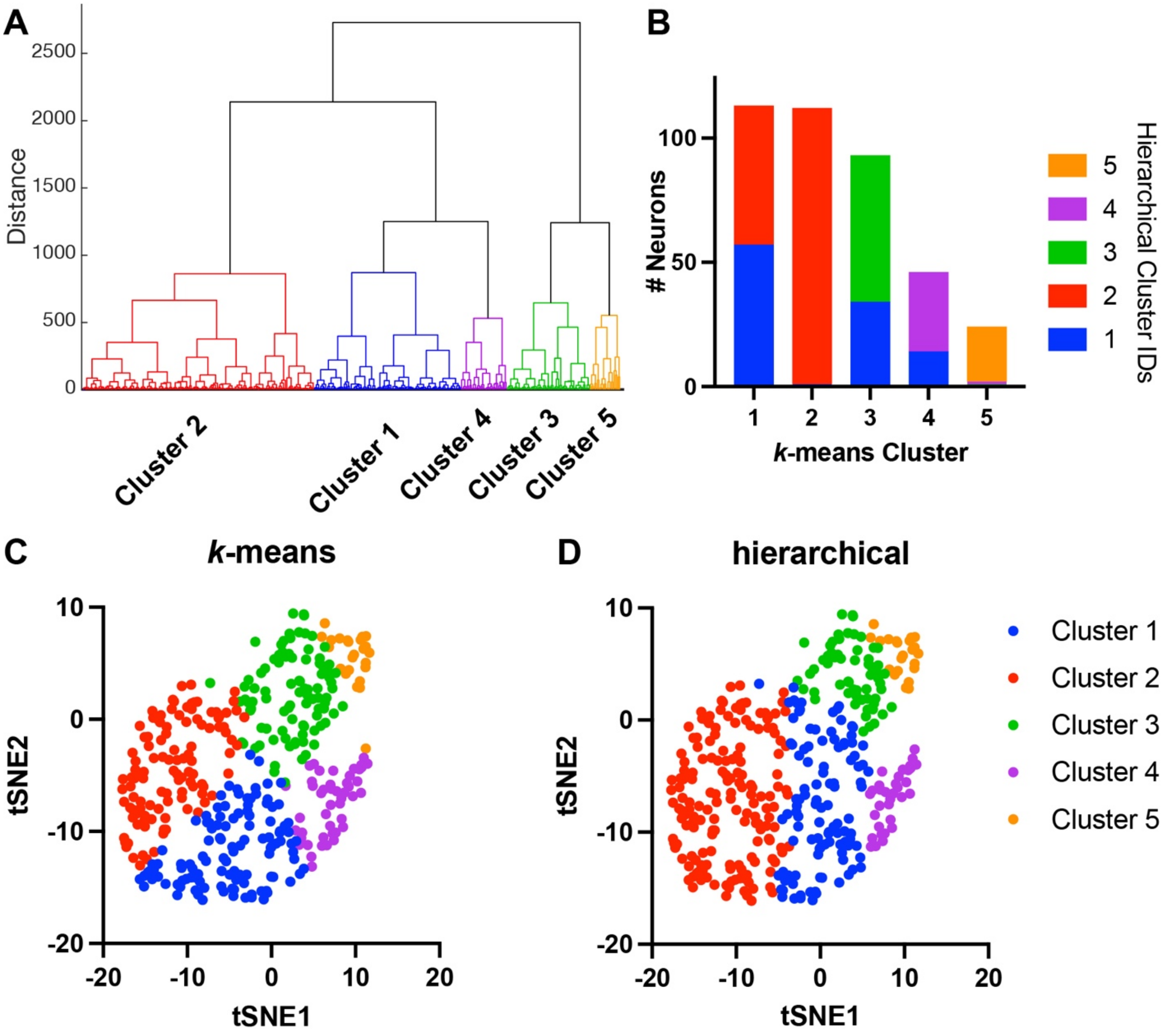
A
Distance
2500
2000
1500
1000
500
0
Cluster 2
Cluster 1
Cluster 4
Cluster 3
Cluster 5
B
### Neurons
100
50
0
1
2
3
4
5
k-means Cluster
Hierarchical Cluster IDs
5
4
3
2
1
C
k-means
D
hierarchical
tSNE2
tSNE1
10
0
-10
-20
-20
-10
0
10
20
Cluster 1
Cluster 2
Cluster 3
Cluster 4
Cluster 5

**Figure S4 – Hierarchical and *k*-means clustering of morphologic features yields similar results.** A) Dendrogram showing relative distance between reconstructed neurons utilizing hierarchical clustering via ward method. B) Quantification of the number of neurons in each *k*-means cluster that had the indicated identity via hierarchical clustering. C-D) Plot of t-Distributed Stochastic Neighbor Embedding (t-SNE) similarities between reconstructed neurons color coded based on cluster identity via *k*-means (C) or hierarchical (D) clustering.

### Figure S5

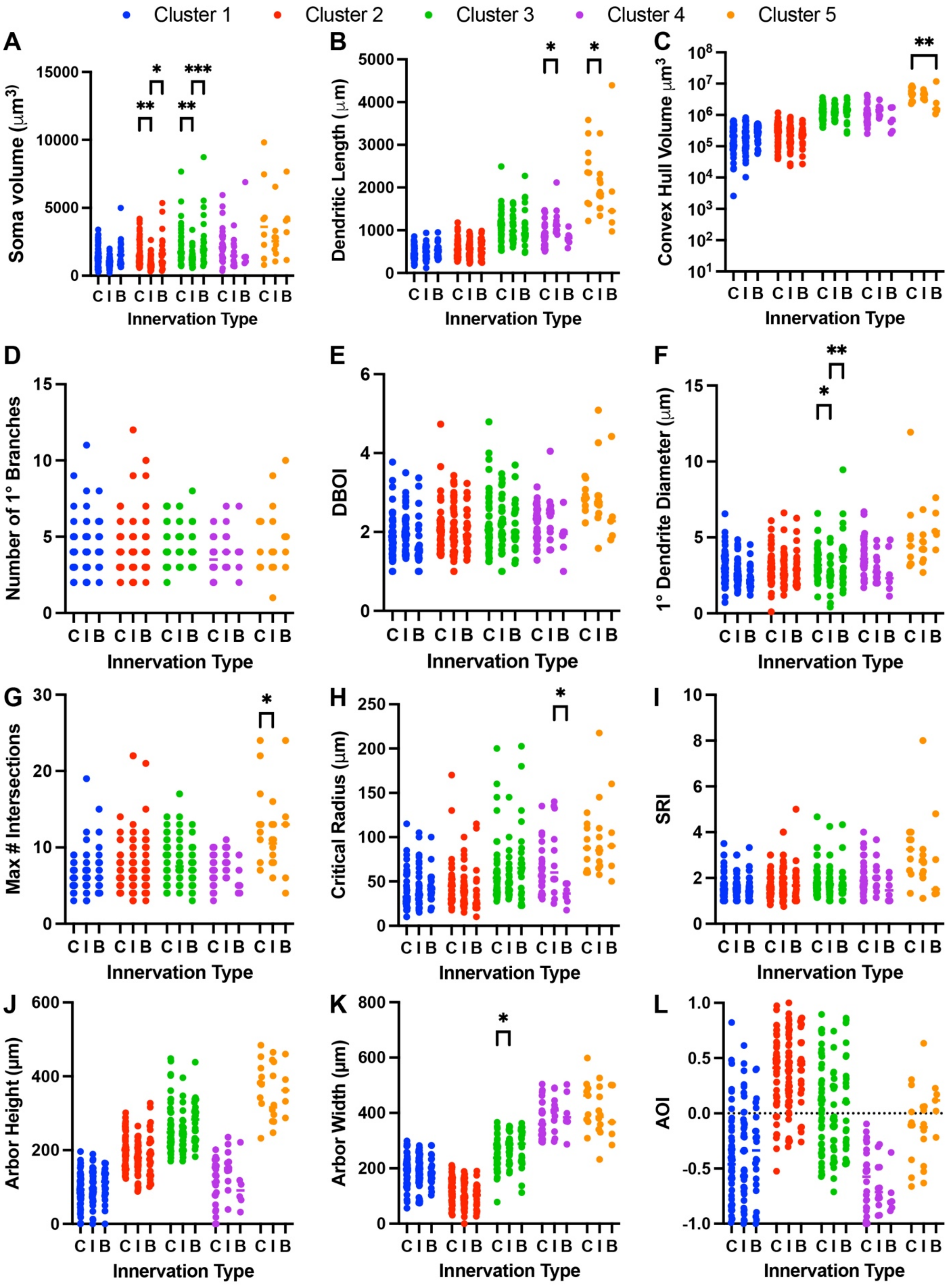

Cluster 1
Cluster 2
Cluster 3
Cluster 4
Cluster 5
A
Soma volume (μm3)
B
Dendritic Length (μm)
C
Convex Hull Volume μm3
D
Number of 1° Branches
E
DBOI
F
1° Dendrite Diameter (μm)
G
Max # Intersections
H
Critical Radius (μm)
I
SRI
J
Arbor Height (μm)
K
Arbor Width (μm)
L
AOI
C I B C I B C I B C I B C I B
Innervation Type

**Figure S5 – Analysis of innervation pattern differences within morphologically-defined subtypes.** A-L) Quantifications of soma volume (A), total dendritic length (B), convex hull volume (C), number of primary dendritic branches (D), dendritic branch order index (DBOI) (E), average primary dendrite diameter (F), the maximum number of intersections (G), critical radius (H), dendritic arbor height (I), dendritic arbor width (J), Sholl ramification index (SRI) (K), and arbor orientation index (AOI) (L) in SC neurons from different clusters identified via *k*-means clustering and innervated by contra-RGCs (*C*), ipsi-RGCs (*I*), or binocularly (*B*). **, $P < 0.05$; **, $P < 0.01$; ***, $P < 0.001$, Kruskal-Wallis test and Dunn's multiple comparisons test*
